# Damage sensing through TLR9 Regulates Inflammatory and Antiviral Responses During Influenza Infection

**DOI:** 10.1101/2024.03.04.583378

**Authors:** Jooyoung Kim, Yifan Yuan, Karen Agaronyan, Amy Zhao, Victoria D Wang, Gayatri Gupta, Heran Essayas, Ayelet Kaminski, John McGovern, Sheeline Yu, Samuel Woo, Chris J. Lee, Shifa Gandhi, Tina Saber, Tayebeh Saleh, Buqu Hu, Ying Sun, Genta Ishikawa, William Bain, John Evankovich, Lujia Chen, HongDuck Yun, Erica L. Herzog, Charles S. Dela Cruz, Changwan Ryu, Lokesh Sharma

**Author notes:** Corresponding authors: Lokesh Sharma at / and Changwan Ryu at. contributed equally as first or senior authors (Jooyoung Kim, Yifan Yuan, Changwan Ryu, Lokesh Sharma).

## Abstract

Host response aimed at eliminating the infecting pathogen, as well as the pathogen itself, can cause tissue injury. Tissue injury leads to the release of a myriad of cellular components including mitochondrial DNA, which the host senses through pattern recognition receptors. How the sensing of tissue injury by the host shapes the anti-pathogen response remains poorly understood. In this study, we utilized mice that are deficient in toll-like receptor-9 (TLR9), which binds to unmethylated CpG DNA sequences such as those present in bacterial and mitochondrial DNA. To avoid direct pathogen sensing by TLR9, we utilized the influenza virus, which lacks ligands for TLR9, to determine how damage sensing by TLR9 contributes to anti-influenza immunity. Our data show that TLR9-mediated sensing of tissue damage promotes an inflammatory response during early infection, driven by the epithelial and myeloid cells. Along with the diminished inflammatory response, the absence of TLR9 led to impaired viral clearance manifested as a higher and prolonged influenza components in myeloid cells including monocytes and macrophages rendering them highly inflammatory. The persistent inflammation driven by infected myeloid cells led to persistent lung injury and impaired recovery in influenza-infected TLR9-/-mice. Further, we show elevated TLR9 activation in the plasma samples of patients with influenza and its association with the disease severity in hospitalized patients, demonstrating its clinical relevance. Overall, we demonstrate an essential role of damage sensing through TLR9 in promoting anti-influenza immunity and inflammatory response.

**Author Summary:** Tissue damage is an inevitable outcome of clinically relevant lung infections, but the host mechanisms for detecting such damage during infection are not well understood. We investigated the role of Toll-like receptor 9 (TLR9) in sensing tissue damage caused by influenza. Since influenza lacks TLR9 ligands, we hypothesized that TLR9 signaling is driven by tissue damage molecules like mitochondrial DNA (mtDNA). Our data indicate that TLR9 reduces early inflammatory lung injury but impairs viral clearance, resulting in extensive immune cell infection, persistent inflammation, and delayed recovery. Myeloid-specific TLR9 deletion ameliorated late-stage inflammatory responses. In humans, influenza-infected individuals exhibited elevated TLR9 activity and mtDNA levels in plasma compared to healthy controls, with higher TLR9 activation potential correlating with severe disease requiring ICU admission. These findings suggest that TLR9-mediated damage sensing triggers both inflammatory tissue injury and viral clearance. These data indicate that TLR9 activity can serve as a crucial biomarker and therapeutic target to limit influenza induced tissue injury.

## Introduction

Toll-like receptors (TLRs) are ubiquitously present innate sensors of specific molecular patterns that are associated with pathogens (PAMPs) or tissue damage (DAMPs)^1^. Sensing pathogens by TLRs leads to an anti-pathogen response while damage sensing is often associated with tissue reparative phenotype^1,2^. However, how the damage-sensing contributes to the anti-pathogen response remains incompletely understood. In a clinically relevant infection, both pathogen-and damage-associated molecular patterns are present simultaneously, making it difficult to dissect the effect of damage sensing on the anti-pathogen response from those generated by pathogen sensing.

Toll-like receptor-9 (TLR9) specifically detects bacterial DNA due to the presence of highly unmethylated CpG domains, enabling it to distinguish it from host DNA^3–5^. In host cells, while nuclear DNA undergoes methylation, mitochondrial DNA (mtDNA) remains highly unmethylated and is detectable by TLR9^6^. During infection-induced tissue injury, mtDNA is often released from dying cells, making mtDNA an important DAMP sensed by TLR9. To understand how the damage signaling contributes to the host defense independent of pathogen sensing, we utilized influenza viral infection, which lacks TLR9 ligands.

Influenza infection is one of the leading causes of infectious disease-induced mortality worldwide with an annual death toll of approximately 500,000^7,8^. Influenza-mediated pandemics have led to some of the biggest loss of human life^9^ and seasonal influenza infections remain a leading cause of morbidity and mortality^7^ despite the widespread availability of vaccines and various antiviral agents. Influenza A often has a zoonotic origin including avian and swine. It is interesting to note that birds lack TLR9^10^ despite influenza being a prominent avian pathogen, emphasizing the dispensable role of TLR9 for influenza sensing.

In light of this knowledge, we sought to determine how tissue damage sensing during influenza infection through TLR9 shapes host response against the influenza virus. In this study, we investigated the role of TLR9 in influenza infection using TLR9-/-mice, meyloid specific TLR9-/-mice, an *ex vivo* cell culture system, human TLR9-/-cells, and peripheral blood samples from patients infected with influenza. Our data show that TLR9 deficiency limits the early inflammatory response and lung injury during influenza infection which comes at the cost of impaired viral clearance. The absence of TLR9 rendered lung cells, especially myeloid cells, susceptible to influenza infection. Furthermore, influenza infection of myeloid cells rendered them highly inflammatory, promoting persistent inflammatory response even after viral clearance. The persistent inflammatory response prolonged the tissue injury and impaired recovery of the host. Deletion of TLR9 in myeloid specific manner preserved early inflamamtory resopnse but ameliorated inflamamtory resposne during recovery phase without affecting viral clearance. Taken together, our findings highlight the key role of damage sensing through TLR9 in promoting pathogen clearance and limiting inflammatory response.

## Results

### TLR9 promotes early inflammatory and injury responses during influenza infection

To understand the role of TLR9-mediated damage sensing in the host response to influenza infection, we infected TLR9-/-and wild type mice (Both B6 background) with 10 PFUs of PR8^11^, a strain that causes significant lung tissue damage^11^. Mice were euthanized at days 4, 7, and 14 post-infection to understand the impact of damage sensing on the time course of infection, including repair. Our data show that WT and TLR9-/-mice had similar inflammatory and injury responses on day 4 (Sup Fig. 1). However, on day 7, relative to WT mice, TLR9 deficient mice demonstrated substantially lower white blood cell (WBC) counts in their bronchoalveolar lavage (BAL, Fig 1A), suggesting a decrease in the overall inflammatory response in the lung. Next, using multicolor flow cytometry (Sup Fig. 2), we found that this reduced WBC count was largely due to decreased macrophage and monocyte recruitment in TLR9-/-mice (Fig. 1B-E). In parallel with this inflammatory response on day 7, TLR9 -/-mice showed a significantly decreased lung injury response as shown by lower BAL total protein content (Fig 1F), RBC counts (Fig 1G), platelet counts (Fig. 1H), and RAGE levels (Fig. 1I).

**Fig. 1.**
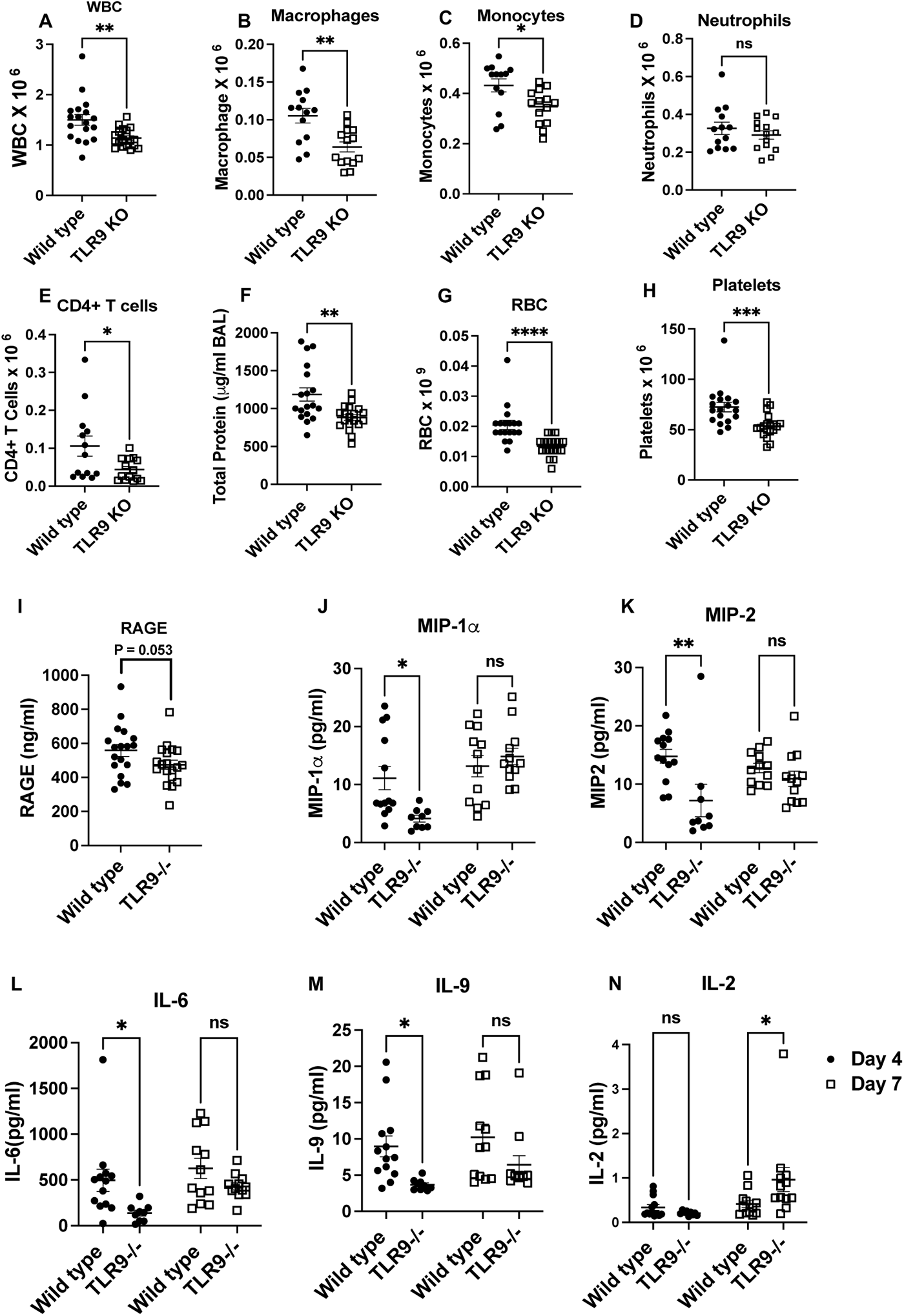
TLR9 signaling promotes early inflammation and injury response during influenza infection: Wild type and TLR9-/-mice were infected with influenza virus by the intranasal route and euthanized on day 7 post infection to harvest broncho-alveolar lavage fluid (BAL). Total white blood cells were counted using a coulter counter (A). The composition of the BAL cells was determined using multicolor flow cytometry. The number of macrophages (B), monocytes (C) neutrophils (D), and CD4+ T cells (E) were obtained. Lung injury was measured by quantifying total protein in the BAL (F) using BCA assay, by measuring the levels of RBCs (G) or platelets (H) in the BAL of mice using a coulter counter and RAGE using ELISA (I). The specific cytokines were measured in the BAL samples using mesoscale discovery multiplex assays (J-N). Two groups were compared with Mann-Whitney analysis, while the cytokine data were analyzed using two-way ANOVA. * = P<0.05, ** = P<0.01, *** = P<0.005, **** = P<0.001, ns = not significant. N = 18 WT and 19 TLR9 KO for A, G and H, pooled from three independent experiments of which flow cytometic analysis was performed for two experiments (N= 13 WT and 14 TLR9 KO) for Fig. B, C, D, E, F, G. Multiplex assays were performed from two independent experiments, N = 13 WT and 9 TLR9 KO for day 4 and N=12 each for WT and TLR9 for day 7.

To further characterize mediators of these ameliorated inflammatory and injury responses, we profiled cytokines associated with inflammatory response (IL-1β, IL-2, IL-4, IL-5, IL-6, IL-9, IL-10, IL-12p70, IL-15, IL-27p28, IL-33, IP-10, KC/GRO, MCP-1, MIP-2α, MIP-2, IFNγ, and TNFα) in the BAL on post-infection days 4 and 7. Our data show that on day 4, relative to WT mice, there was a reduction in the macrophage/monocyte cytokines such as MIP-1α and MIP-2, along with a decrease in IL-6 and IL-9; both mice exhibited similar levels of these soluble mediators on day 7 (Fig. 1J-M). In contrast, we observed a significant increase in IL-2 levels on day 7 (Fig. 1N). We observed no significant changes in the other cytokine concentrations between WT and TLR9 -/-mice on day 4 or 7. These findings suggest that injury sensing through TLR9 mediates both inflammatory and injury responses in the lung in the early stages of influenza infection.

### Damage signaling through TLR9 promotes viral clearance independent of interferon response

Having found the contribution of TLR9-mediated damage sensing in lung inflammation and injury response to influenza, we then endeavored to identify the role of TLR9 in viral clearance. On day 7, relative to their WT counterparts, TLR9 deficient mice exhibited higher viral loads by rt-qPCR (Fig. 2A) and by Western blotting of influenza NS1 protein (Fig. 2B and C). However, we did not find any difference in replicating virus as measured by TCID50 assay (Fig. 2D), indicating a role of TLR9 signaling in promoting non-productive viral replication. The elevated viral protein signature persisted even at day 10 post-infection before getting cleared by day 14, a time point when the replicating virus was cleared from both genotypes (Sup Fig. 3). Interestingly, this increased viral load was not related to an impairment of type I and type III interferon response. We observed similar early interferon response at early time point of day 4 (Sup Fig. 1D and E). On day 7, we observed similar levels of IFNα (Fig. 2E), but an enhanced type 1 interferon response in the BAL of TLR9 deficient mice (Fig 2F). The elevated levels of type I interferons in the BAL on day 7 are likely a consequence of elevated viral fragments present in immune cells. This is of significance since the interferon levels are elevated despite the decreased inflammatory cell recruitment on day 7 (Fig. 1A-1E). Further, despite increased viral load, histological analysis did not show increasaed lung pathology in TLR9 KO mice on day 7 (Sup Fig. 1F).

**Fig. 2.**
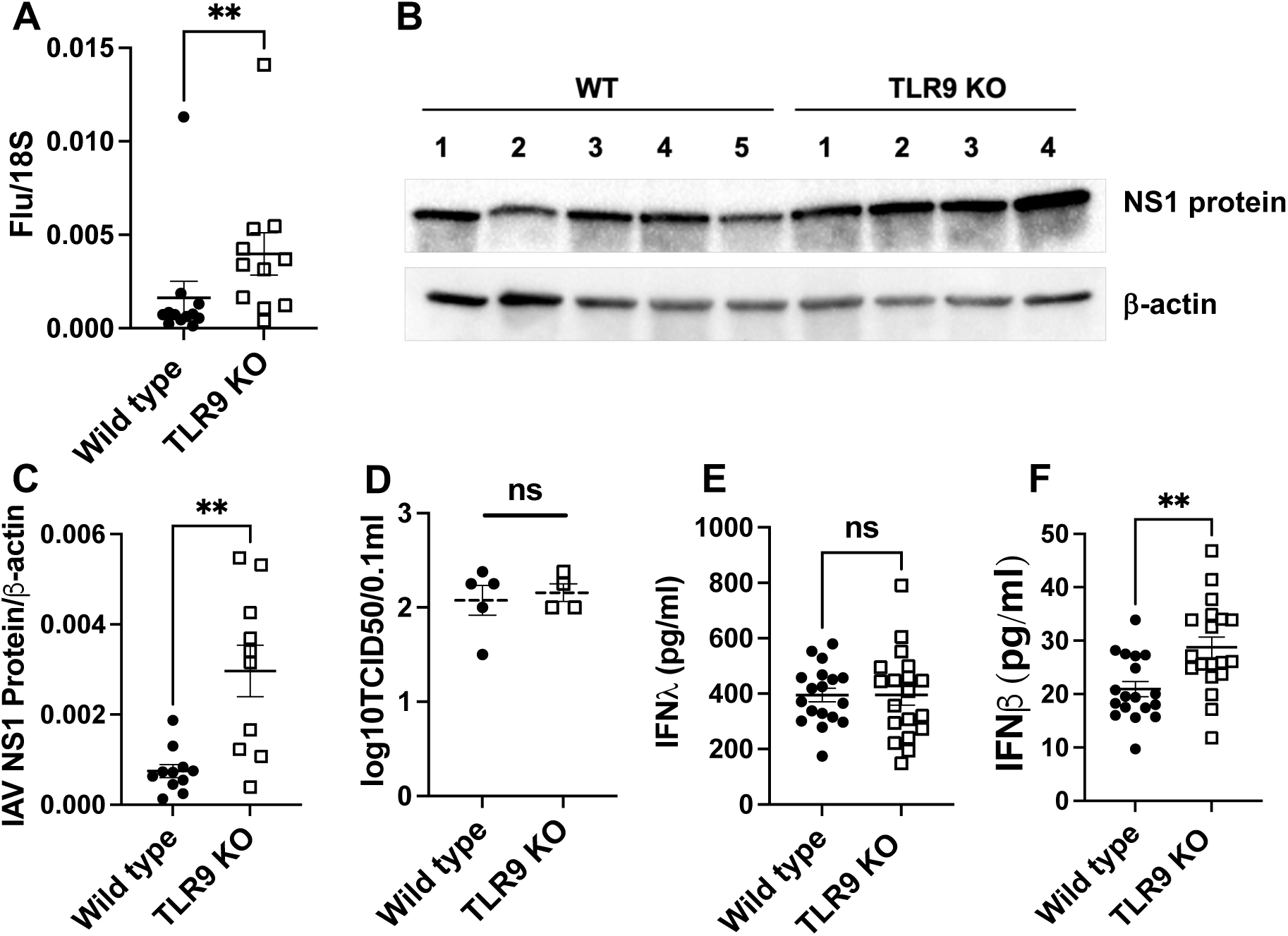
TLR9 signaling limits non-productive viral growth in influenza infection: Viral load was measured in the lung tissue of wild type and TLR9-/-mice on day 7 post-infection using the qPCR of viral matrix gene (A) where 18S was used as housekeeping control gene. The viral load was confirmed using Western blotting of the lung tissue using antibodies against the N protein of influenza. Western blot analysis (B) and densitometric analysis (C) are shown. Beta-actin was used as housekeeping control. Productive viral replication was measured using TCID50 using MDCK cell culture (D). Levels of interferon β (E) or interferon λ (F) were measured using sandwich ELISA. Data were analyzed using Mann-Whitney analysis. * = P<0.05, ** = P<0.01, ***, ns = not significant. Each dot represents a mouse, N = 12 WT and 11 KO for A and C pooled from two experiments and 18 and 19 from three experiments.

### TLR9 deficiency leads to infection of innate immune cells

To gain mechanistic insights into the contribution of damage sensing through TLR9 in nonproductive viral clearance during influenza infection, we performed single-cell RNA-sequencing of lung tissue. Lung samples were obtained on day 4 post-infection to determine cell-specific differences in early infection and antiviral responses. Our single-cell approach yielded significant populations of both immune cells including macrophages, monocytes, neutrophils, T cells, B cells, basophils, dendritic cells, and epithelial cells including ciliated, AT1, AT2, goblet, and secretory epithelial cells (Fig. 3A and Sup Fig. 4A). The cellular specificity was determined based on the canonical gene signatures^12,13^ as indicated in Fig. 3B and Sup Fig. 4B. We determined the presence of viral infection in these cells by influenza transcripts to understand the differences in the cellular targets of influenza infection in TLR9-/-lungs. A minimal infection was observed across the immune cell types in wild-type mice on day 4 post-infection. In contrast, TLR9-/-immune cells demonstrated significantly more infections which were spread across all the immune cells including innate immune cells such as neutrophils, monocytes, and macrophages (Fig. 3C and D). A distinct monocyte population, that expanded during viral infection, had high influenza infectivity denoted as iMon_high infection (Fig. 3A). These cells expanded during influenza infection in both wild type and TLR9-/-mice, however, a more robust expansion was observed in TLR9-/-mice (Sup Fig. 5). The increased viral burden in TLR9-/-deficient cells were not limited to only immune cell compartment, we found elevated infection in the epithelial compartment as well (Sup Fig. 4 C & D).

**Fig. 3.**
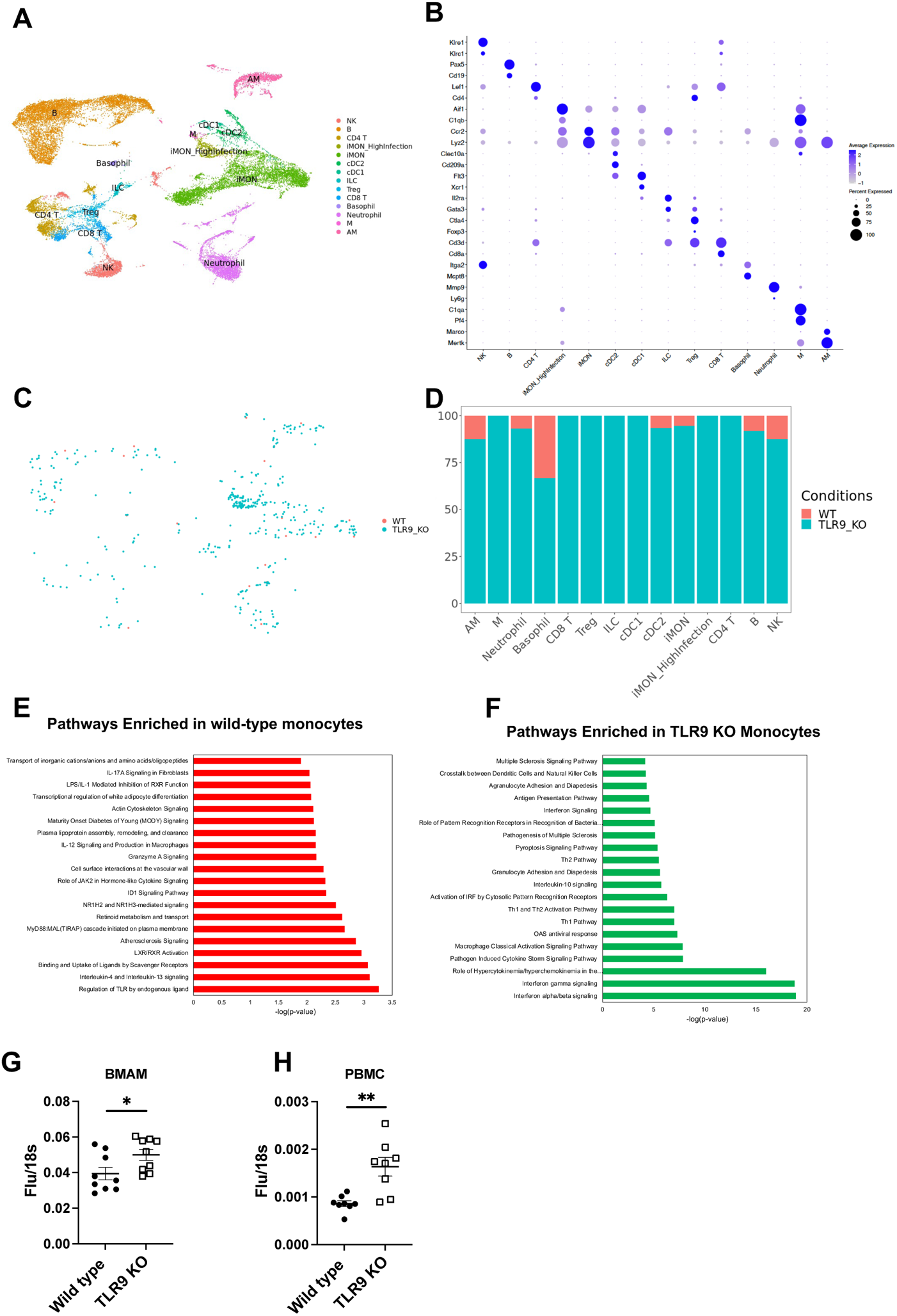
TLR9 deficiency renders immune cells susceptible to influenza infection: Uniform Manifold Approximation and Projection (UMAPs) plots of immune cells present in the lungs based on the known gene expression markers (A). iMON_Highinfection was the subset of inflammatory monocytes that were infected with the influenza virus and were exclusively present in the TLR9-/-mice. Among the immune cells, the influenza-infected cells were identified using influenza RNA expression in the cell and separated by the genotype (B). The infection status was quantified based on the genotype to compare wild type vs TLR9-/-cells (C). Top pathways enriched in the wild type (D) or TLR9-/- (E) inflammatory monocytes are demonstrated. Bone marrow-derived alveolar macrophages were infected with influenza virus (MOI of 1) for 24 hours and the RNA was isolated to perform qPCR of influenza matrix protein (F). TLR9-/-human peripheral blood monocyte cells were generated and infected with the influenza virus similarly to measure the viral load (G). Data are pooled from 3 independent experiments. Data were analyzed using Mann-Whitney analysis. * = P<0.05, ** = P<0.01, ***.

To better understand the impact of damage sensing on the host antiviral response, we investigated differentially expressed genes in wild-type vs TLR9-/-immune cells. TLR9-/-monocytes upregulated chemokines and cytokines and associated receptors including Ccl3, Ccl4, Ccr5, IL-15, and IL-18. In contrast, we observed an increase in Bcl2, Cd36, and fibronectin-1 in the wild-type monocytes (Sup Fig. 6A). To determine whether these high expression of cytokines and chemokines interact with epithelial cells to have functional consequences at later time points, we generated Circos plots to understand cell-cell communications. Using iMON as signal senders (ligands) and epithelial cells such as Secretory, AT2, and Ciliated cells as signal receivers (receptors), we identified several ligand-receptor pairs such as Ccr2, Ccr10, Il1r, Il-12rb, Il15ra with their multiple chemokine and cytokine receptors (Sup Fig. 6B). Diff-Connectome analysis demonstrate ligand-receptor interactions are upregulated in TLR9_KO cells.

To understand pathways associated with these gene changes, we performed IPA analysis based on differentially expressed gene signatures in inflammatory monocytes. Wild-type monocytes demonstrated enrichment of tissue reparative pathways such as “interleukin 4 and interleukin-13 signaling” and “Binding and uptake of ligands by scavenger receptor”. Further, as expected “Regulation of TLR by endogenous ligand” was the top upregulated pathway in the wild type monocytes (Fig. 3E). In contrast, the type I interferon pathway was the most significantly enriched in TLR9-/-monocytes (Fig. 3F). This observation was in agreement with our data demonstrating elevated type I interferon levels in TLR9-/-mice *in vivo* (Fig. 2F). Interestingly, these enriched pathways were unique to monocytes and not shared by epithelial cells such as ciliated epithelium (Sup Fig. 4 E & F)). To understand whether TLR9-/-deficiency leads to increased susceptibility to influenza infection in immune cells, we directly infected wild-type and TLR9-/-alveolar macrophages *ex vivo*. Our data show that TLR9-/-alveolar macrophages had inherent defects in the viral control as higher viral replication was observed in these cells (Fig. 3G). A similar finding was observed in human cells TLR9-deficient peripheral blood mononuclear cells (PBMCs). Infection of these cells demonstrated impaired viral control in PBMCs with TLR9 deficiency (Fig. 3H).

### Immune cells concurrently upregulate influenza and inflammatory gene signature

To understand how extensive infection of TLR9-/-immune cells affect their inflammatory response, we determined both the influenza response and inflammatory gene signature in these cells. For the measurement of inflammatory gene signature in the immune cells, we used “hallmark_inflammatory_response” module that includes 197 mouse genes (GSEA M5932) to show that elevated inflammatory response is present in the immune cells, especially that of myeloid origins such as monocytes, macrophages and neutrophils, which was further upregulated by influenza infection (Fig. 4A). Similarly, we explored the “influenza response” signature using 36 genes to demonstrate that influenza upregulated this gene signature in both wild-type and to a higher extent in TLR9-/-myeloid cells (Fig. 4B). This was supported by a signficant upregulation of STAT1 and NFkB pathways in the immune cells in absence of TLR9 (Sup Fig. 7A and B). Furthermore, we performed a crossover analysis between inflammatory and influenza gene signatures to show a significant overlap, especially in myeloid cells including monocytes, macrophages, and neutrophils in both these signatures (Fig. 4C). Next, we investigated whether similar elevation of STAT1 and NFkB was observed in epithelial cell compartment. Our data show that similar to immune cells, TLR9 KO epithelial cells upregulated STAT1 compared to wild type cells (Sup Fig. 7C), a likely consequence of elevated viral burden and subsequent interferon responses in these cells. However, in contrast to immune cells, we observed a signficant downregulation of NFkB pathway in TLR9 KO epithelial cells (Sup Fig. 7D), potentially contributing to dampened early cytokine and inflamamtory response in TLR9 KO mice during influenza infection (Fig. 1).

**Fig. 4.**
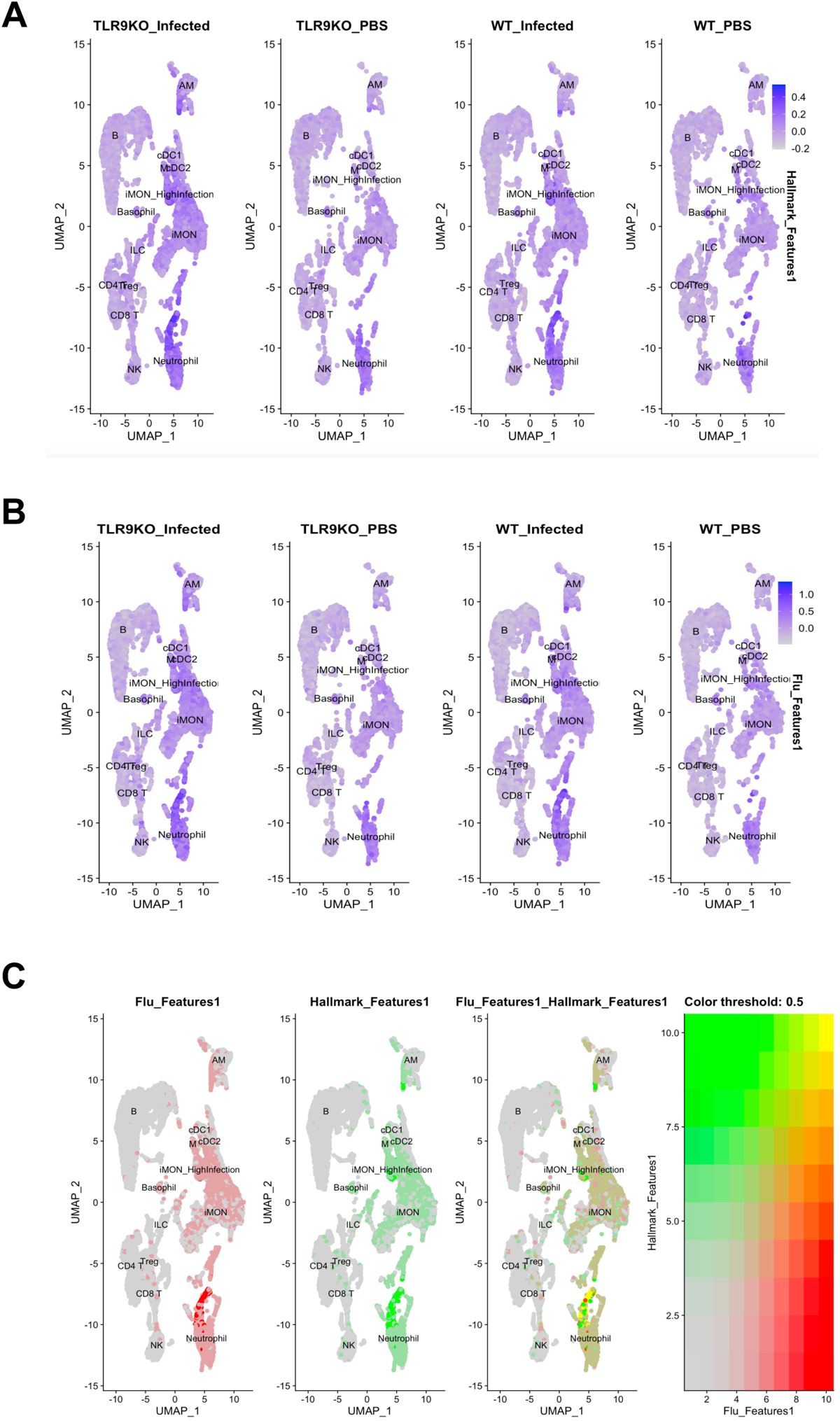
Influenza infection simultaneously upregulates both influenza and inflammatory response in immune cells: The expression of the “hallmark_inflammatory_response” module that includes 197 mouse genes (GSEA M5932) in the immune cells (A) and influenza response (B). The crossover between “hallmark_inflammatory_response” and “influenza response” in the immune cells (C).

**Fig. 5.**
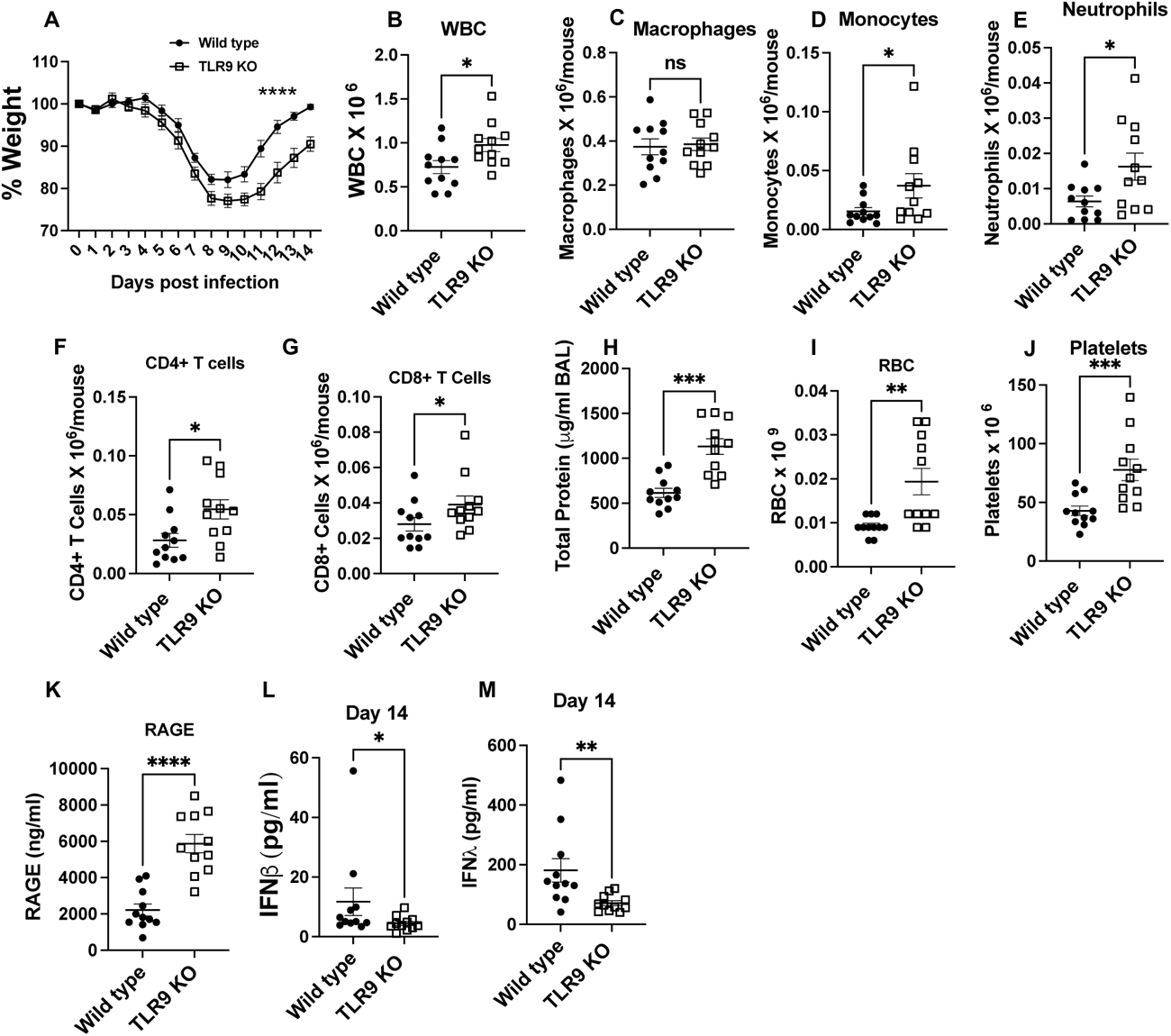
TLR9-/-mice have impaired recovery during influenza infection: Wild type and TLR9-/-mice were infected with influenza virus and body weight was measured every day for 14 days (A). Mice were euthanized on day 14 to harvest broncho-alveolar lavage. Total WBCs in the BAL were counted using coulter counter (B). The number of macrophages (C), monocytes (D), neutrophils (E), CD4+ (F), and CD8+ T (G) cells was determined using multi-color flow cytometry. Lung injury was measured by measuring total protein content (H), RBC counts (I), platelets (J) and RAGE (K) in the BAL. Levels of interferon b (L) and interferon l (M) were measured in the BAL obtained on day 14 post infection. Data are pooled from two independent experiments. Weight loss curves were analyzed using two-way ANOVA with Šídák’s multiple comparisons test. Two groups were compared with Mann-Whitney analysis, * = P<0.05, ** = P<0.01, *** = P<0.005, **** = P<0.001, ns = not significant, N=11 each group.

Next, to find the consequences of these upregulated inflammatory pathway in immune cells, we investigated cell death pathways. Our data show that influenza infection upregulated multiple cell death pathways including apoptosis, necroptosis, and pyroptosis pathways in immune cells, implicating excessive infection in immune cell death in absence of TLR9 (Sup Fig. 8).

To understand how human myeloid cells respond to infection, we performed time course of inflammatory response in human PBMC cell line. Our data show that while TLR9 deficient cells had ameliorated inflammatory responses during early time points (2 or 6 hours), the inflammatory response in TLR9-/-cells increased dramatically as the infection progresses (Sup Fig. 6C). These data support the hypothesis that TLR9 deficiency ameliorates the early inflammation through dampened inflamamtory response by epithelial and immune cells, however, this advantage wanes due to excessive replication of influenza in absence of TLR9, especially in immune cells.

### Elevated viral load impaired recovery in TLR9-/-mice

We observed a reduction in early inflammation and injury response in TLR9-/-mice (Fig. 1), however, it came at a cost of impaired viral control (Fig. 2 and 3). Further, we observed an elevated inflammatory response in the myeloid cells that had influenza gene signatures (Fig. 4). We next sought to determine how this impaired viral clearance contributes to the recovery in mice from influenza infection. We followed wild-type and TLR9-/-mice for up to 14 days post influenza infection. Our data show that TLR9-/-mice had significantly impaired recovery manifested as delayed body weight gains (Fig. 5A). The impaired recovery in TLR9-/-mice was associated with a persistent inflammatory response in the lung, manifested as elevated immune cell numbers in the BAL (Fig. 5B). Significantly elevated levels of monocytes and neutrophils persisted in the TLR9-/-mice at this time, while the number of macrophages in the BAL were similar at this time point (Fig. 5C-E). We also saw an increased recruitment of CD4+ and CD8+ T cells in the airspaces of TLR9-/-mice on day 14 (Fig. 5F and G). This persistent inflammatory response was also associated with increased tissue injury manifested as elevated total protein content (Fig. 5H), along with increased RBC (5I), platelet counts (5J), and RAGE in the BAL (5K). However, at this time, despite elevated inflammatory cell recruitment in these mice, we observed a significant reduction in both type I and type III interferon response (Fig. 5L and M) indicating that TLR9 signaling controls interferon signaling in the absence of a replicating virus (Fig. 3C). Despite elevated inflamamtory response, we did not observe an elevated fibrotic response at this time point assessed by trichrome staining of lung tissues (Sup Fig. 9A). Lack of increase fibrosis in TLR9 KO mice despite elevated inflamamtory resopnse is a likely consequence of dynamic changes in inflamamtory injury during the course of influenza infection.

To determine whether this impaired viral recovery is due to elevated viral load or TLR9 have independent role in promoting host recovery in viral infection, we stimulated mice with TLR9 agonist on day 8 post viral infection. Our data show that treatment with TLR9 agonist ODN2006 at a time point when most of the viral load is cleared, impaired host recovery (Supplemental Fig. 9B) indicating that impaired recovery in TLR9-/-mice is likely due to persistent presence of viral infection. This was further supported by experiments utilizing antiviral drug oseltamivir in mice, which blocked the impaired recovery phenotype in TLR9-/-mice (Sup Fig. 9C), emphasizing a role of impaired viral control in recovery.

The contribution of impaired viral control was further supported by utilization of myeloid specific TLR9 knockout mice. In contrast to the global deficiency, myeloid specific deletion of TLR9 did not lead to impaired viral control (Sup Fig. 10A). However, it ameliorated inflammatory response during the recovery phase on day 14 (Sup Fig. 10B), further supporting the notion that elevated viral load in global TLR9 KO contributes to the impaired recovery. The ameliorated inflammatory response in LysM Cre+TLR9-/-mice was mainly attributed to the myeloid cells including macrophages and monocytes (Sup Fig. 10C). These data demonstrate that TLR9 signaling acts on multiple cells to shape host response to influenza infections.

### Increased TLR9 ligands in patients with influenza infection

Finally, to determine the clinical relevance of our findings, we sought to measure the levels of TLR9 ligand mtDNA in the clinical samples of patients with influenza infection. We recruited influenza-infected hospitalized patients at Yale New Haven Hospital and obtained the age and sex-matched control samples. The detailed demographics are presented in Fig. 6A. Our data show that compared to plasma samples from healthy controls, those from individuals infected with influenza displayed significantly higher TLR9 activation as observed with a human reporter cell line (Fig. 6B). To further confirm the TLR9 activity of plasma samples, we measured the levels of mtDNA in these samples (Fig. 6C). To investigate whether elevated TLR9 signaing or mtDNA concentrations are relevant to the clinical disease, we compared these levels between those who were admitted to the general medical ward versus those who required a higher level of care in the medical intensive care unit (MICU). Our data show that the overall TLR9 activation by patient plasma was significantly elevated in those who were admitted to MICU as compared to those who did not (Fig. 6D). We did not observe similar effect with mtDNA, indicating that receptor activation is more relevant than ligand levels. To further investigate this biology, we measured the TLR9 activity of mouse serum during the time course of influenza infection using a mouse TLR9 reporter cell line. Our data showed that a signficant elevation in TLR9 activity was observed on day 7 (peak inflammation) (Sup Fig. 11), indicating a potential correlation between mouse model and human disease. Together these data indicate that circulating TLR9 activity serves as a biomaker for influenza infection and associated disease severity.

**Fig. 6.**
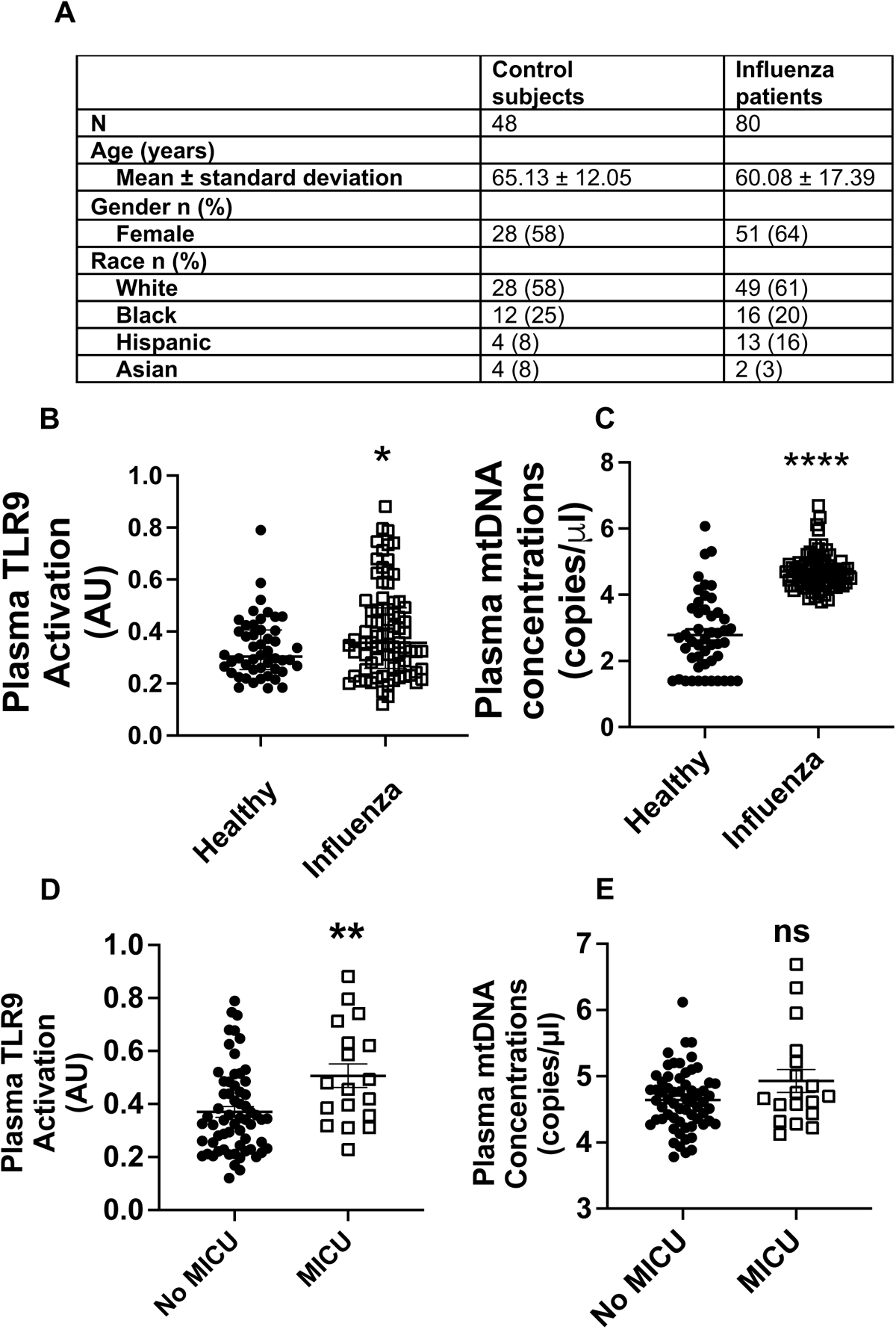
Human influenza infection upregulates TLR9 ligands in circulating blood: Plasma samples were obtained from patients with influenza infection and control subjects without any respiratory infections. The total number of subjects in each group and demographics are shown in Fig. 6A. The overall TLR9 activity of plasma was measured using reporter cell line (B). The levels of mtDNA were measured using qPCR-based assay (C). Plasma TLR9 activity (D) and mtDNA levels (E) were compared between those admitted to medical intensive care unit (MICU) and those remained hospitalized without the need of MICU, as marker of disease severity. *, P<0.05, ** = P<0.01, **** = P<0.001 using Mann-Whitney test.

## Discussion

In this study, we identified an essential role of damage sensing through TLR9 on host defense mechanisms during influenza infection. During an infection, the host balances its response to eliminate the pathogen (host resistance) while limiting the collateral tissue damage (host tolerance). Host defense and host tolerance are distinct strategies that a host employs to survive an infection^14,15^. However, how the host fine-tunes the inflammatory response during an infection and how the tissue damage shapes the antipathogen response still needs to be completely understood. In this study, we demonstrate that the absence of damage sensing through TLR9 deficiency improved the host tolerance to influenza infection at the early stage of the infection by limiting early inflammatory response by epithelial and immune cells (Fig. 1). However, this absence of damage sensing impaired the host defense, manifested as elevated and prolonged presence of viral remnants (Fig. 2), which ultimately leads to persistent injury and inflammatory response (Fig. 5), compromising the disease recovery.

Other damage-sensing receptors such as TLR4, which senses oxidized phospholipids, contribute to pathogenic inflammation and tissue injury, including in influenza infection^16,17^. However, unlike our observations for TLR9, the absence of TLR4 did not impair the host defense against influenza infection, leading to overall better outcomes^16^. This highlights the complexity of damage sensing where specific DAMP molecules contribute differentially to the host resistance and host tolerance. A complete understanding of damage sensing through various receptors might provide us with therapeutic targets that don’t have potential deleterious effects on host immunity.

Our data also shed critical insights into how TLR9 signaling contributes to the host ability to limit viral replications, especially those by immune cells where nonproductive viral replication takes place. TLR9 deficiency led to extensive infection of immune cells such as macrophages, inflammatory monocytes, and neutrophils. The increase in the viral load in the immune cells appeared to be independent of conventional antiviral pathways mediated by type I interferons^18^. We observed a dichotomy between wild-type and TLR9-/-immune cells in their response to influenza infection where wild-type cells upregulated genes that were required for homeostasis and tissue regeneration such as “binding and uptake of scavenger receptors”^19^ and “retinoid metabolism”^20^. In contrast, TLR9-/-cells upregulated the genes that were associated with an inflammatory milieu such as “hypercytokinemia/hyperchemokinemia response” or “pathogen-induced cytokine storm”. It is critical to note, that these changes, especially in long-living cells such as monocytes and macrophages could drive the persistence of inflammatory milieu, as observed in TLR9-/-mice. These data shed light on the clinical manifestations of influenza infections where pathological inflammation persists even after the clearance of actively replicating virus and contributes to the lethality. This hypothesis is further supported by prior studies in influenza virus where presence of viral remnants contributes to the persistent inflammation at the site of infection much after the replicating virus is cleared^21^. Although it is possible that extensive infection in the myeloid cells may not lead to effective replication and release of infectious virus, it may be sufficient to promote persistent inflammation and delayed recovery, as observed in TLR9-/-mice.

There are limited studies that have been performed to investigate the role of damage-sensing through TLR9 in influenza infections. Previous studies have indicated that stimulation of TLR9 along with TLR2/6 can provide robust protection against influenza infection by promoting viral clearane^22^, which seemed to be independent of the immune cell compartment^23^ and dependent on the cell’s ability to generate reactive oxygen species^24^. The potential interaction of TLR9 with TLR2 has also been demonstrated by their synergistic role in detecting certain strains^25^ of herpes simplex virus-1 through plasmacytoid dendritic cells. Similarly, the absence of TLR9 renders the host susceptible to certain viruses such as cytomegalovirus^26^, however, initial evidence indicated that it does not affect the clearance of influenza infection^27^ but controls post-viral bacterial infections. The role of TLR9 in recognizing the bacterial DNA and mounting a protective host response during pulmonary Streptococcus infection has been demonstrated previously^28^. However, these studies either boost TLR9 signaling before the tissue damage^22–24^ or utilize a secondary bacterial infection that can be detected by TLR9 through bacterial DNA^27^. Thus, our study is the first to demonstrate the indispensable role of damage signaling through TLR9 in limiting viral replication and persistence.

To ensure the translational aspects of our findings, we utilized human peripheral blood mononuclear cells that were deficient in TLR9 to demonstrate the role of TLR9 in controlling influenza infection in immune cells. Our human data also corroborated mouse data demonstrating that increased viral load in the TLR9-/-PBMCs led to elevated inflammatory response. Further, we demonstrated the enhanced activity of TLR9 among patients who were hospitalized with influenza infection, especially those admitted to the MICU (Fig. 6). TLR9 activation was similiarly observed in mouse model during peak disease on day 7 (Sup Fig. 11), indicating its potential to serve as a biomaker for influenza disease severity. We also observed a dichotomy between mtDNA and overall TLR9 activity, which can be explained by other DAMPs present during the disease state. Prior studies have shown that other DAMP molecule HMGB1 has been known to modulate the TLR9 activating potential of mtDNA^29^.

Overall, our study demonstrated that damage sensing through TLR9 plays a key role in the anti-influenza immune response. This anti-influenza response appeared to be independent of type I interferon signaling, a well-established pathway downstream of TLR9 signaling.

## Material and Methods

### Mouse model of influenza infection

Wild type and TLR9-/-mice were obtained from Jackson laboratory on C57 B/6 background and bred in house in the same room at the Yale University. TLR9 flox mice were generated and provided by Dr. Mark Shlomchik at Yale University (currently at University of Pittsburgh). Mice were infected with influenza virus (H1N1, PR8) by intranasal route as previously described. Mice were anesthetized using ketamine/xylazine solution and an infection inoculum of 10 PFUs in 50 μl of PBS was administered by intranasal route. The control infection mice received the same amount of PBS. ODN2006 was administered by intranasal route. All the animal work performed was approved by Institutional Animal Care and Use Committee at Yale School of Medicine (Protocol #20044).

### Human Subjects

All the human research was approved by the Institutional Review Board at Yale School of Medicine with HIC# 0901004619. All the patient samples and health volunteers were recruited after obtaining written or verbal (in presence of a witness) informed consent.

### *In vitro* culture of murine alveolar macrophage and human peripheral blood monocytes

Alveolar macrophages were cultured from bone marrow cultures of wild type and TLR9-/-mice in the presence of granulocyte-macrophage colony-stimulating factor, TGFβ, and the PPARγ activator as described recently^30^. The expression of various alveolar markers was confirmed by qPCR (data available at https://figshare.com/articles/figure/Characterization_of_bone_marrow_derived_alveolar_macrophages_/26253473). The TLR9 knockout (homozygous) human peripheral blood mononuclear cells were generated by Synthego via CRISPR-Cas9 technology. Guided RNA sequence used for TLR9 knockdown was: AGGCUGGUGACAUUGCCACG. The knockdown was validated by PCR using the following primers: FOR Primer (5’-3’): CTGCCTTCCTACCCTGTGAG and REV Primer (5’-3’): GAGTGACAGGTGGGTGAGGT.

### Cell counting and flow cytometry

White blood cell counts (WBC), red blood cells (RBCs), and platelets were counted using a Coulter counter. Flow cytometry was performed with multicolor flow cytometry protocols similar to those published previously.

### Western Blot

Western blot was performed as described previously by our lab^31^. At indicated time points, mice were euthanized and gathered whole lungs. Tissues were placed in a lysis buffer and homogenized with PRO25D Digital homogenizer. Tissue lysates were cleared of debris by centrifugation (15min, 3000g) and mixed with 2x Laemmli Sample Buffer. Equal amounts of protein (30 μg) were resolved by SDS-PAGE using 4-20% Mini-PROTEAN^®^ TGX Stain-Free Precast™ Gels gels and transferred onto Trans-Blot^®^ Turbo™ membranes (Bio-Rad). Membranes were blocked with 5% non-fat milk in TBS-T for 1 hour at room temperature and probed with primary antibodies against (Influenza A virus NS1) overnight at 4°C. After incubation with HRP-conjugated secondary antibodies, protein bands were visualized using an enhanced chemiluminescence detection kit (Thermo Fisher Scientific).

### qPCR

Total RNA was extracted from whole lungs from IAV non-infected / infected mice and human PBMCs using (RNeasy mini kit, Qiagen). RNA quantity and purity were assessed using a spectrophotometer (NanoDrop). Subsequently, 1μg of total RNA was reverse transcribed into cDNA using a high-capacity cDNA reverse transcription kit (iScript cDNA Synthesis Kit, Bio-Rad) according to the provided protocol. Specific targeting primers were designed using Primer3 software and validated for efficiency and specificity. Primer sequences were as follows: Flu forward; AAGACCAATCCTGTCACCTCTGA, Flu reverse; CAAAGCGTCTACGCTGCAGTCC, hTNF-α forward; CTCTTCTGCCTGCTGCACTTTG, hTNF-α reverse ; ATGGGCTACAGGCTTGTCACTC, hIL-6 forward; AGACAGCCACTCACCTCTTCAG and hIL-6 reverse; TTCTGCCAGTGCCTCTTTGCTG. Real-time qPCR was performed using a (Bio-Rad CFX384 Real-Time 384-well PCR qPCR Detection System Pred CFX Opus – AV) with SsoAdvanced Universal SYBR^®^ Green Supermix (Bio-Rad) under the following cycling conditions: initial denaturation at 95°C for 10 minutes, followed by 40 cycles of denaturation at 95°C for 15 seconds and annealing/extension at 60°C for 1 minute.

### Detection of mitochondrial DNA

DNA was extracted from plasma using the QiaAMP DNA Mini-Kit (Qiagen) as previously described^32^. Briefly, 200 µl of plasma sample was used to elute 100 µl of DNA. The presence of human mtDNA in each sample of DNA was assayed by qPCR for the MT-ATP6 gene, using primers for MT-ATP6 (ThermoFisher) and probes (SsoAdvanced Universal Probes Supermix, Bio-Rad Laboratories). The ViiA7 Real-Time PCR System (ThermoFisher) was employed with the following cycling conditions: incubation at 95°C for 10 minutes; 40 amplification cycles at 95°C for 10 seconds, 60°C for 30 seconds, and 72°C for 1 second; and cooling at 40°C for 30 seconds. The number of MT-ATP6 copies per microliter was determined based on a standard curve developed using serial dilutions of a commercially available DNA plasmid (OriGene) with complementary DNA sequences for human MT-ATP6.

### ELISA and Multiplex

BAL from IAV non-infected / infected mice were assayed for cytokine levels using commercially available sandwich ELISA kits for RAGE, IFN-β and IFN-α(R&D Systems). Multiplex kits from MSD were used as per the manufacturer’s instructions.

### Statistics

Data were analyzed using either student’s t-test or one-way ANOVA with Tukey’s multiple comparison test. Statistical significance was assumed when p values were <0.05.

### 10x scRNAseq Library Preparation, and Sequencing

Lung samples across different conditions were digested with collagenase type 4 (2mg/ml) and DNase (20U/ml) in DMEM with 10% FBS to prepare single-cell suspension for an expected cell recovery population of 10,000 cells per lane. scRNAseq libraries were generated with the Chromium Single Cell 3’ assay (10x Genomics) and were sequenced on the NovaSeq platform per library in Yale Center for Genomics Analysis. Downstream processing was conducted with Cellranger 3.0.2 with the default parameters.

RNA-Seq Data Filtration, Normalization, Scaling, Integration, and Clustering: Aligned data was processed using Seurat v3.1.2 under R environment. Default parameters were used for each function unless otherwise specified ^33^. After creating Seurat objects for each sample, we filtered on the percent mitochondrial genes to exclude damaged and dying cells and on the number of genes and transcripts to remove debris and doublets. All samples after filtration were integrated by the IntegrateData function in Seurat. The gene counts in the integrated object were then normalized to the total counts and multiplied by a 10,000 scaling factor. We used Variance Stabilizing Transformation (VST) to select highly variable genes. The top 200 genes within the selected genes were retained for downstream analysis. Variable genes were scaled using Seurat’s ScaleData function and subsequently were used for principal component analysis (PCA) to generate clusters. The numbers of clusters produced using PCA analysis are controlled by a resolution parameter with higher values giving more clusters and by the number of PCs. All cells across different samples were assigned into two-dimensional Uniform Manifold Approximation and Projection (UMAPs).

To sub-cluster immune cells and epithelial cells, we first used subset function in Seurat to select cell clusters that are positive for Ptprc, and Epcam while negative for Col1a1, and Cdh5 to create a new lung immune cell or epithelial cell Seurat objects, respectively. We then re-normalized, re-scaled, and re-clustered these objects for downstream analysis.

Cluster Identification by Differentially Expressed Genes (DEGs): The FindAllMarkers function in Seurat was applied to each sample to identify differentially expressed genes for each cluster relative to all the other clusters in the object. We only chose genes that were expressed in at least 10% of cells in one of these clusters to ensure the quality of genes. These marker lists were used to apply cluster labels based on top defining genes of canonical cell types in the literature ^12,13^.

To identify viral reads in the sequencing data, the influenza A genomic reference was created with CellRanger mkref. The sample fastq files were then aligned to this reference genome using CellRanger count. Barcodes corresponding to reads that aligned with the influenza virus were considered “infected.” This information was transferred to the single-cell object’s metadata.

After pre-processing and quality control, two gene module scores were calculated: (1) influenza-related genes, and (2) hallmark inflammatory genes. Influenza gene lists were generated by searching the top 30 genes associated with the keywords “influenza” in Geneshot Pubmed. The hallmark inflammatory genes were obtained from the GSEA gene list ‘hallmark_inflammatory_responsè (GSEA M5932). For each cell, the composite gene module score was calculated. These two scores were then plotted, split by condition. The overlap between gene lists (1) and (2) was plotted using the Seurat FeaturePlot blend option.

Gene lists for:

Influenza signature: ACE2, ANP32A, BST2, CXCL10, DDX58, IFITM1, IFITM3, IFNA1, IFNB1, IFNG, IFNL1, IL10, IL1B, IL6, MAVS, MBL2, MX1, NFKB1, NLRP3, RAB11A, SFTPD, SOCS1, TGFB1, TLR2, TLR3, TLR4, TLR7, TMPRSS2, TNF, TP53.

STAT1 pathway:

Stat1, Ifng, Stat3, Il6, Tnf, Il1b, Jak2, Jak1, Ifna1, Cd4, Il10, Il4, Ifnb1, Irf1, Stat6, Actb, Cd8a, Gapdh, Akt1, Stat5a, Stat5b, Socs1, Stat2, Il17a, Tgfb1, Pxdn, Pxdnl, Socs3, Ccl2, Cd274

NFkB pathway:

Nfkb1, Il6, Tnf, Il1b, Nfkbia, Actb, Akt1, Tlr4, Casp3, Gapdh, Il10, Ifng, Tnfsf11, Jun, Alb, Ptgs2, Ccl2, Il1a, Tgfb1, Cd4, Tp53, Ins, Pxdn, Pxdnl, Bcl2, Mapk3, Ikbkb, Mmp9, Stat3, Aif1

Pyroptosis:

Casp1, Il1b, Gsdmd, Il18, Nlrp3, Gapdh, Actb, Il6, Tnf, Casp3, Tlr4, Alb, Gsdma, Pxdn, Pxdnl, Gsdme, Casp8, Nlrc4, Aim2, Cck, Anxa5, Ifng, Pycard, Il10, Mlkl, Gpt, Il1A, Itgam, Nlrp1, Ripk3

Apoptosis:

Casp3, Bcl2, Anxa5, Tp53, Akt1, Actb, Gapdh, Tnf, Pxdn, Pxdnl, Il1b, Il6, Cycs, Alb, Bax, Casp9, Dntt, Pik3ca, Mtor, Pik3cg, Pik3cb, Pik3cd, Casp8, Bcl2L1, Ifng, Cck, Tgfb1, Parp1, Ins, Cd4

Necroptosis:

Mlkl, Ripk3, Ripk1, Tnf, Casp3, Casp8, Gapdh, Il6, Actb, Il1b,Pcsk1, Alb, Anxa5, Casp1, Fadd, Pxdn, Pxdnl, Tnfrsf1a, Ifng, Tlr4, Akt1, Casp9, Cck, Il18, Ins, Nfkb1, Aif1, Cd4, Dntt, Gpt

## Methods for Circos plots and Diff-Connectome

Cell-cell communication analyses were performed using the R package Connectome. Average expression levels of ligands and receptors per cell type were computed across experimental groups. Connectomes were constructed for each experimental group, containing unfiltered lists of edges linking ligand-expressing cells to receptor-expressing cells.

To compare TLR9 KO and WT infected mice, log-fold changes in expression were evaluated for both ligand and receptor sides of all edges using *FindMarkers*. A perturbation score was calculated to facilitate differential edge plotting, reflecting the log-fold change and incorporating directional information from ligands and receptors. Edges meeting criteria (at least 10% involvement of sending and receiving clusters, adjusted p-value < 0.05 via Wilcoxon rank sum test) were visualized using *CircosPlot*.

## Data Availability

Raw data used to generate the figures of the manuscript has been provided as an excel file (Supporting Table 1). Single cell RNA seq data has been deposited to Gene Expression Omnibus under the accession number GSE271505.

---

**Supplemental Fig. 1.**
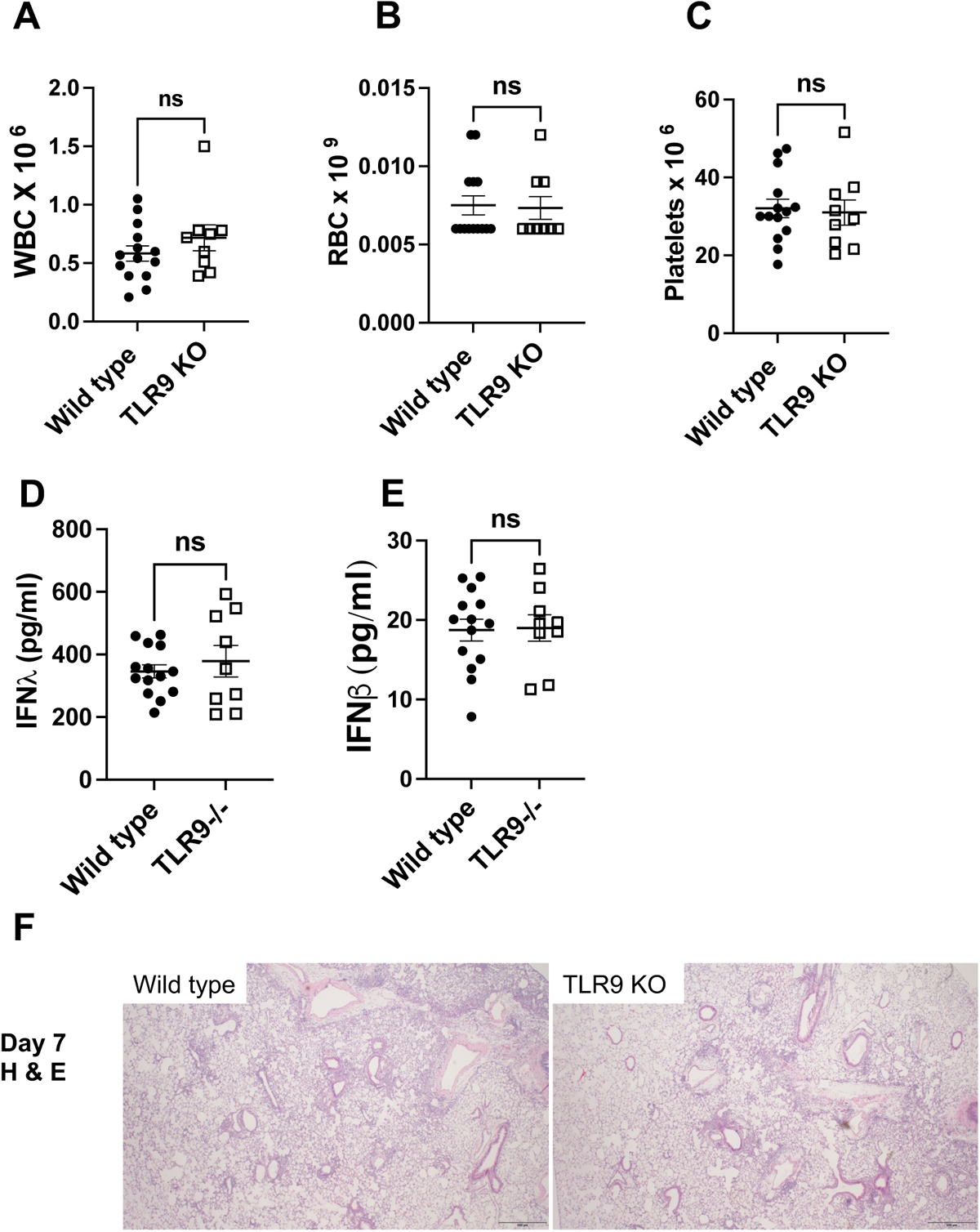
Levels of total WBC, RBC, and platelets were counted using a Coulter counter on day 4 post infection in wild type and TLR9-/-mice (A-C). Interferon levels were measured using ELISA (D and E) Lung sections were stained with hematoxylin and eosin on day 7. Sup Fig. 1F shows representative pictures on day 7.

**Supplemental Fig. 2.**
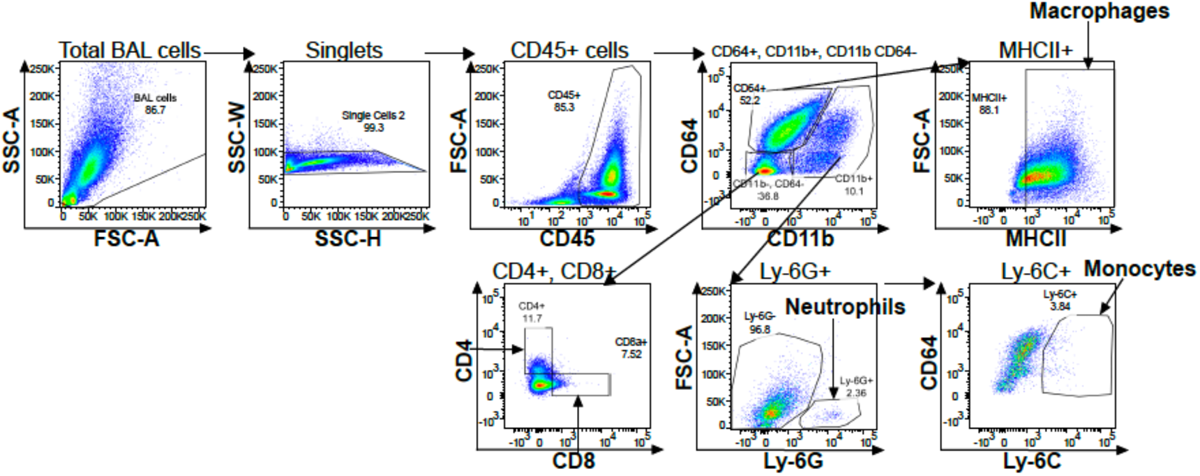
Flow strategy to detect immune cell populations in the BAL.

**Supplemental Fig. 3.**
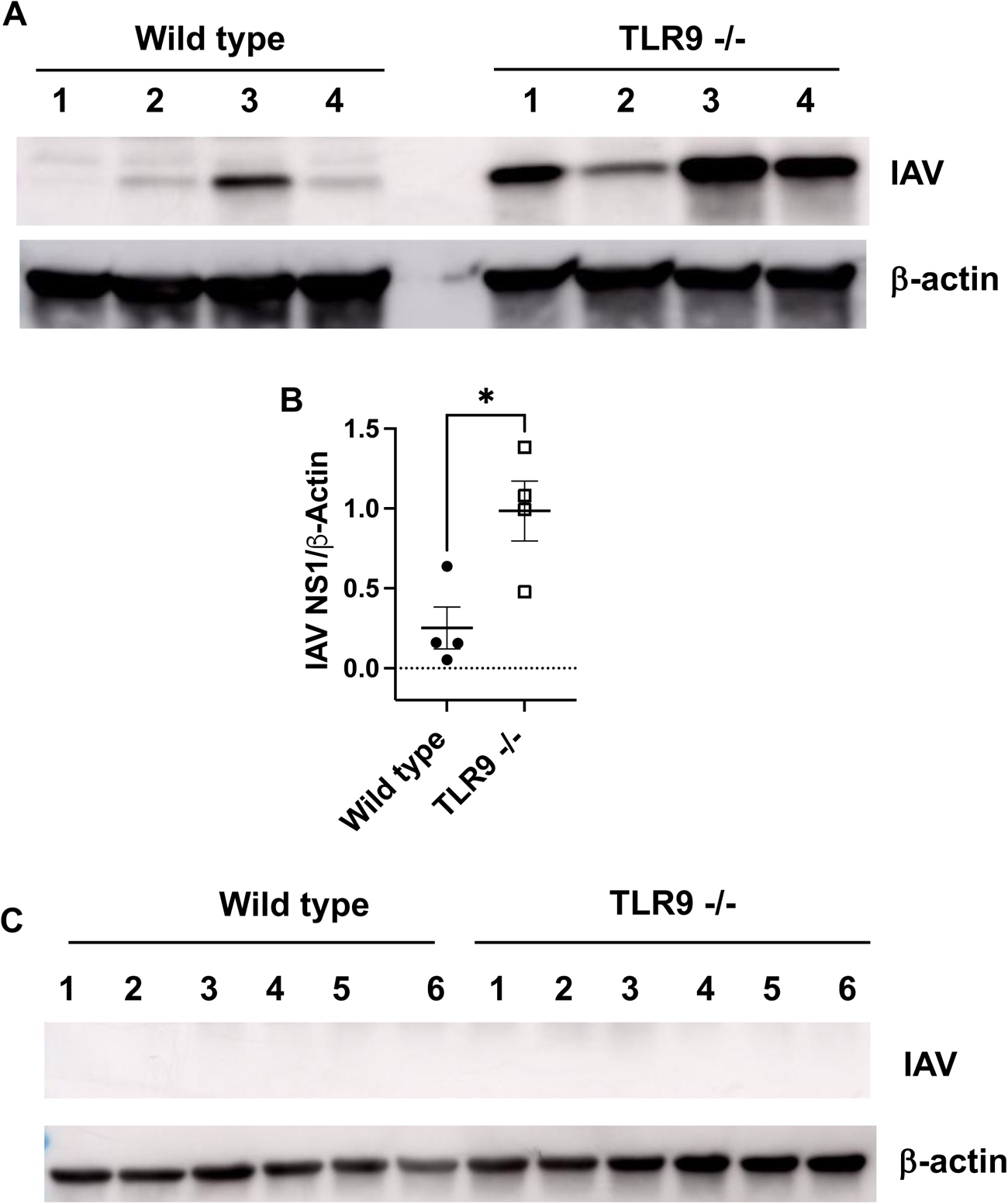
Western blot analysis of influenza NS1 protein on day 10 post-infection (A and B) and day 14 post infection (C).

**Supplemental Fig. 4.**
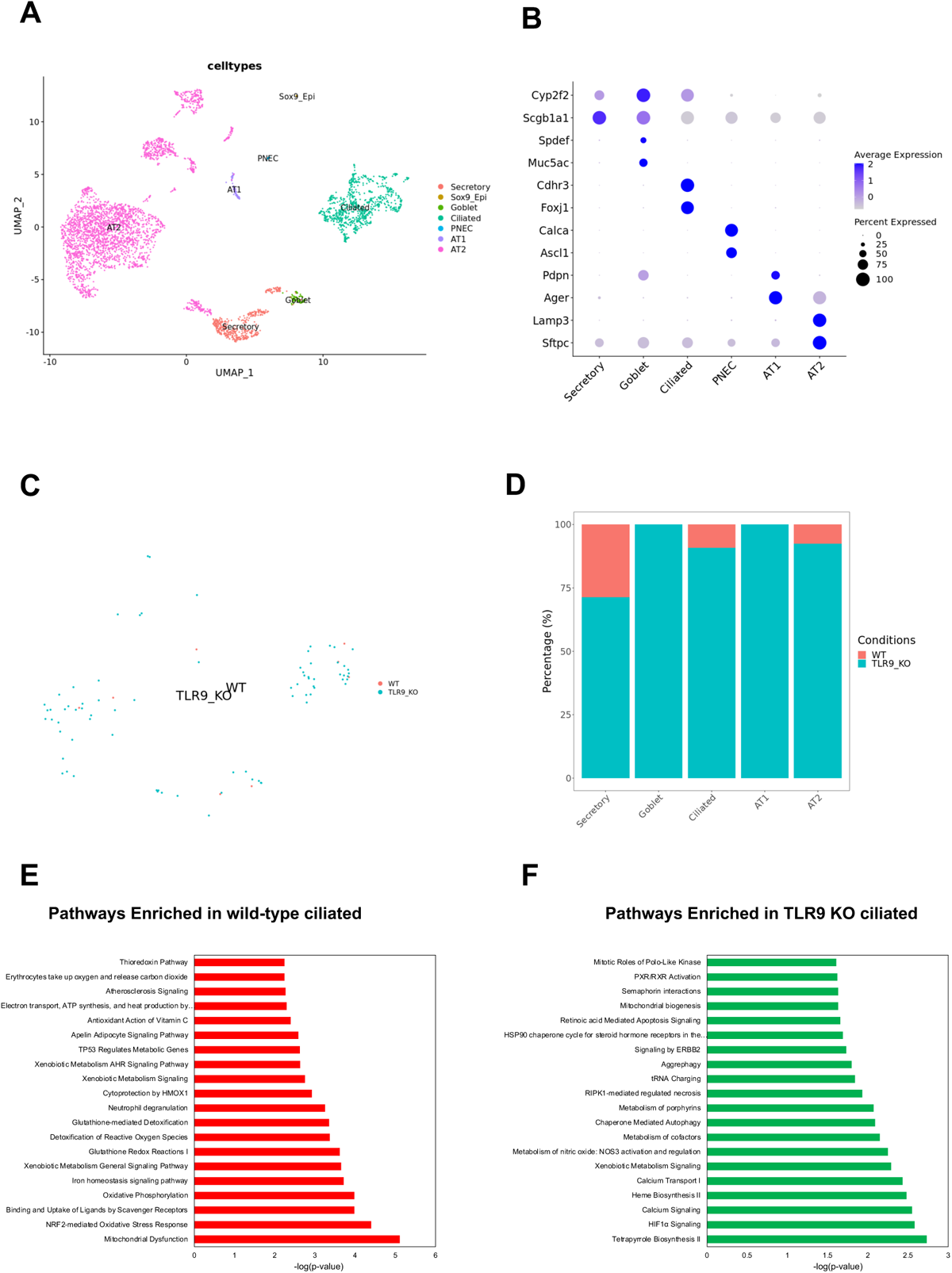
UMAP of epithelial cells in the lung (A) and markers to identify specific cell types (B). The identity of infected cells and associated genotype (C) and quantification (D). IPA pathway analysis showing enriched pathway in wild type and TLR9-/-ciliated epithelium (E and F).

**Supplemental Fig. 5.**
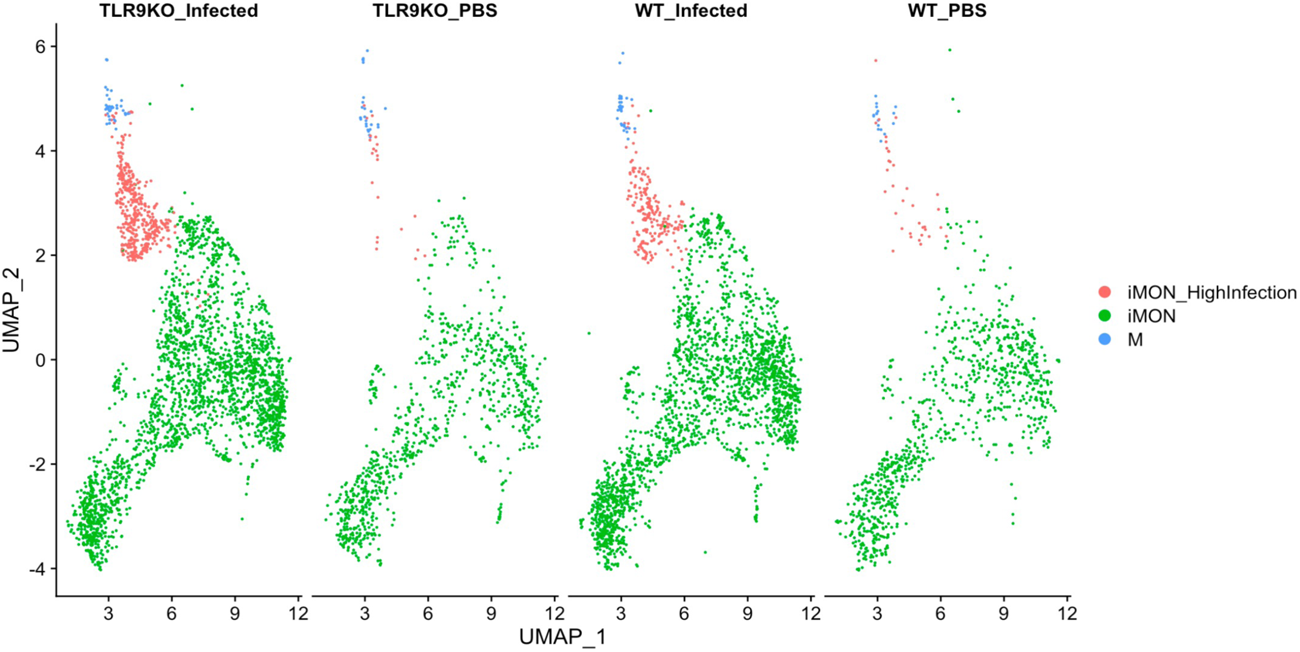
UMAP showing an expansion of iMON_Highinfection in infected mice, especially in TLR9-/-lungs.

**Supplemental Fig. 6.**
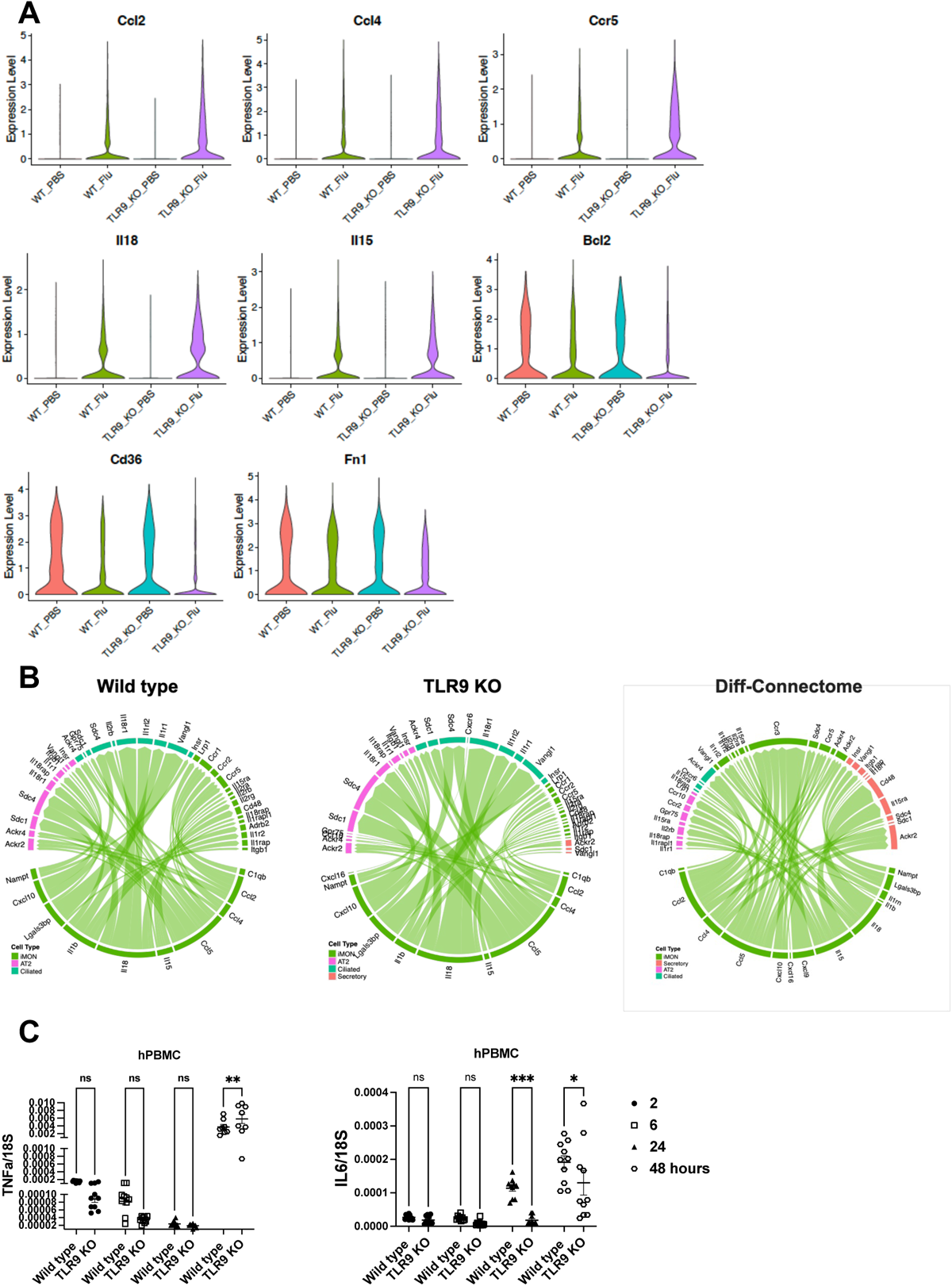
Violin plots showing the expression levels of specific gene (A). Cell-cell communication analyses were performed using the R package Connectome. Average expression levels of ligands and receptors per cell type were computed across experimental groups. Connectomes were constructed for each experimental group, containing unfiltered lists of edges linking ligand-expressing cells to receptor-expressing cells. Diff-connectome was generated to demonstrate the upregulated receptor-ligand interactions in TLR9 KO cells (B). Human PBMC cell lines were infected with influenza (MOI of 1) and cytokine expressions were measured over the time course using qPCR (C). *, P < 0.05, **, P<0.01, and ***, P < 0.005 using two-way ANOVA.

**Supplemental Fig. 7.**
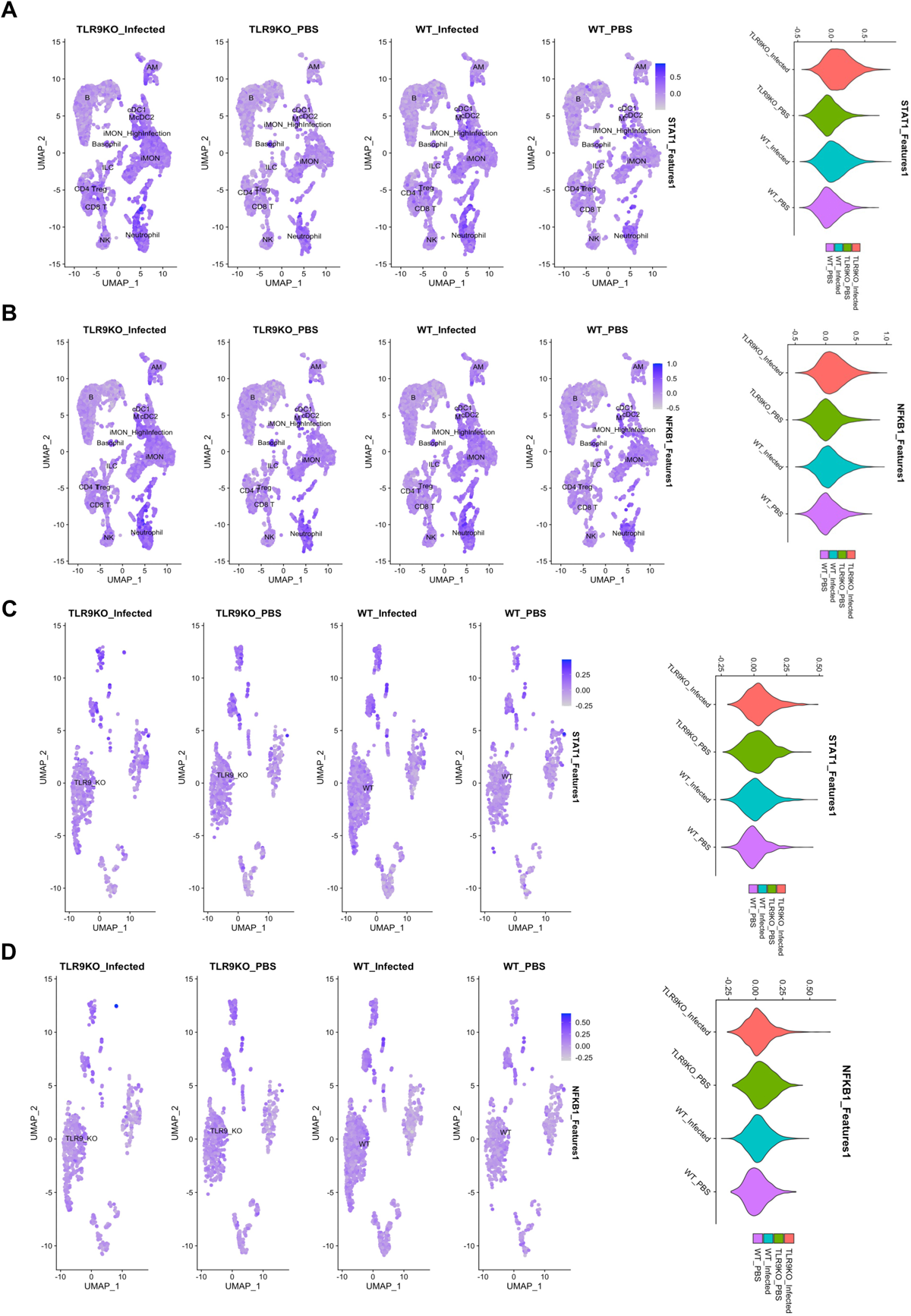
STAT1 and NFkB pathway signature in immune (A and B) and epithelial cells (C and D) along with respective quantifications on the right.

**Supplemental Fig. 8.**
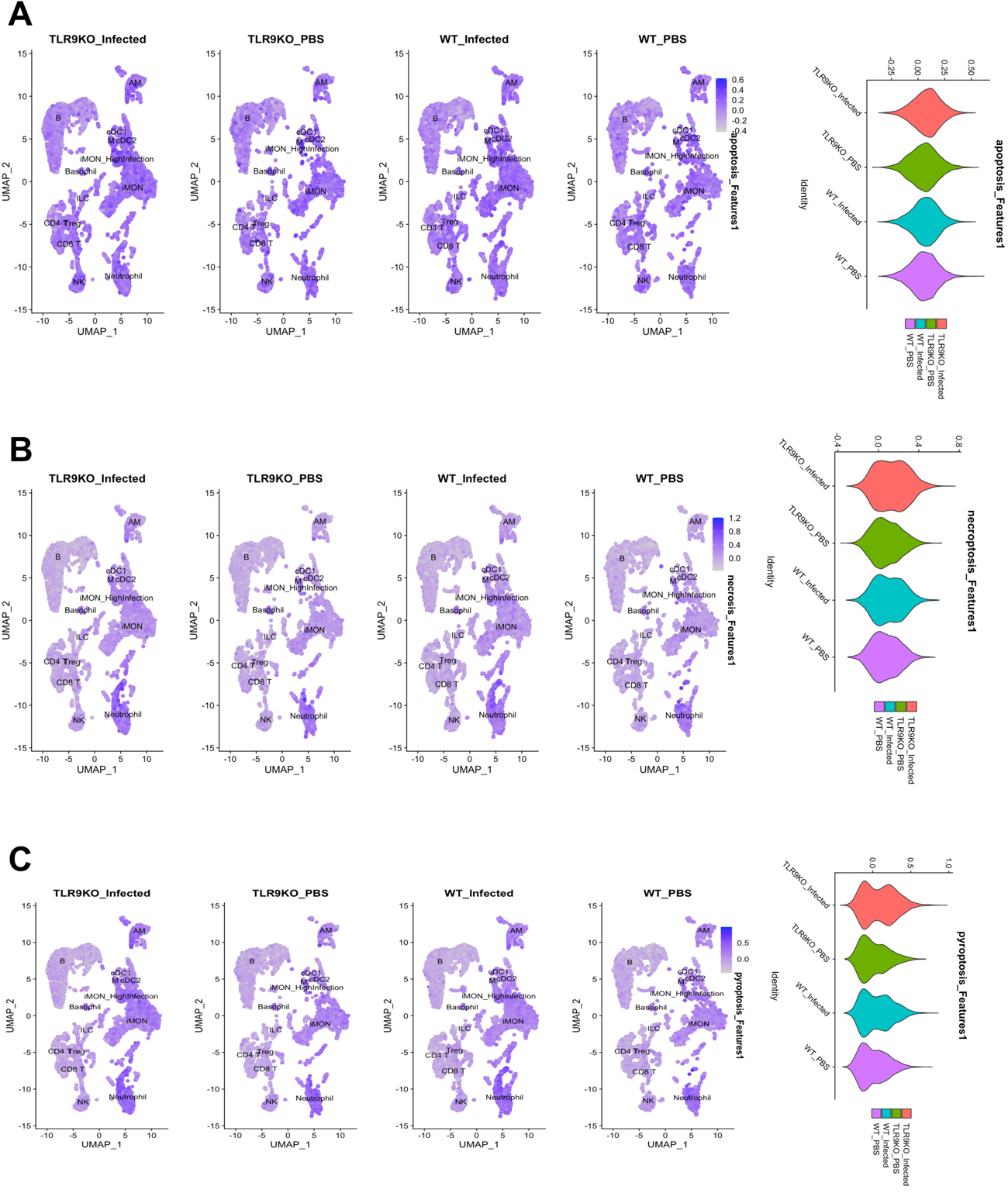
Apoptosis (A), necroptosis (B), and pyroptosis (C) pathway signatures in immune cells along with respective quantifications on the right.

**Supplemental Fig. 9.**
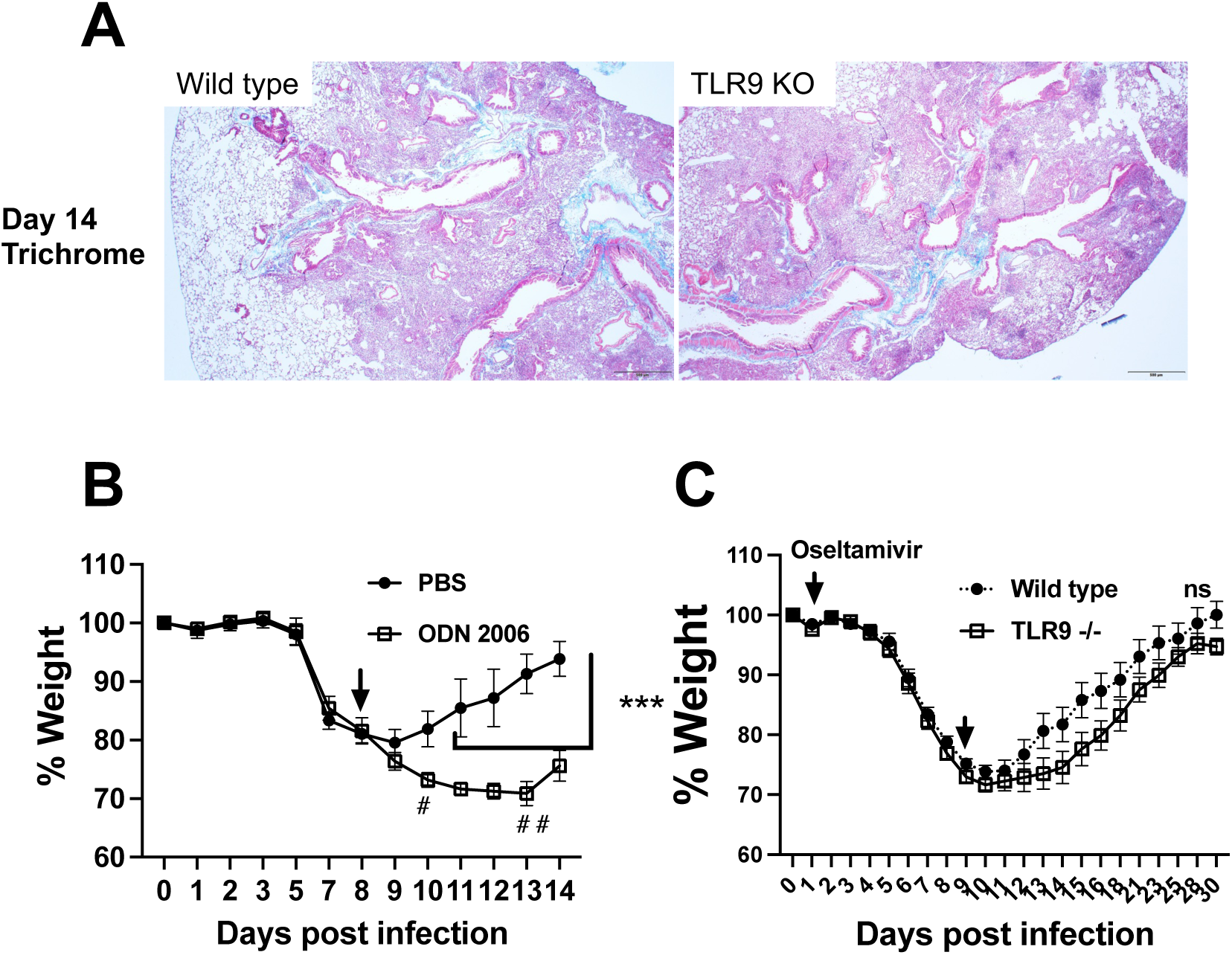
TLR9 stimulation post viral clearance impairs recovery: Histological analysis of lung tissue sections that were stained with trichrome staining to assess fibrosis (blue color) (A). Wild type mice were infected with influenza virus and randomized into two groups to either receive TLR9 agonist ODN2006 or PBS on day 8 by intranasal route. Body weights were measured every day. # indicates mortality in the ODN2006 on the indicated day (B). Wild type and TLR9-/-mice were infected with 4x dose (40 PFUs) of influenza virus and treated with oseltamivir every day (30 mg/kg) starting on day 1 post infection and until day 9 post infection. Body weights were measured (C). N=4-5 each group in A and 9-10 each group in B. ***, P<0.005 using two-way ANOVA with Šídák’s multiple comparisons test, n= non-significant, # indicates one death on that day.

**Supplemental Fig. 10.**
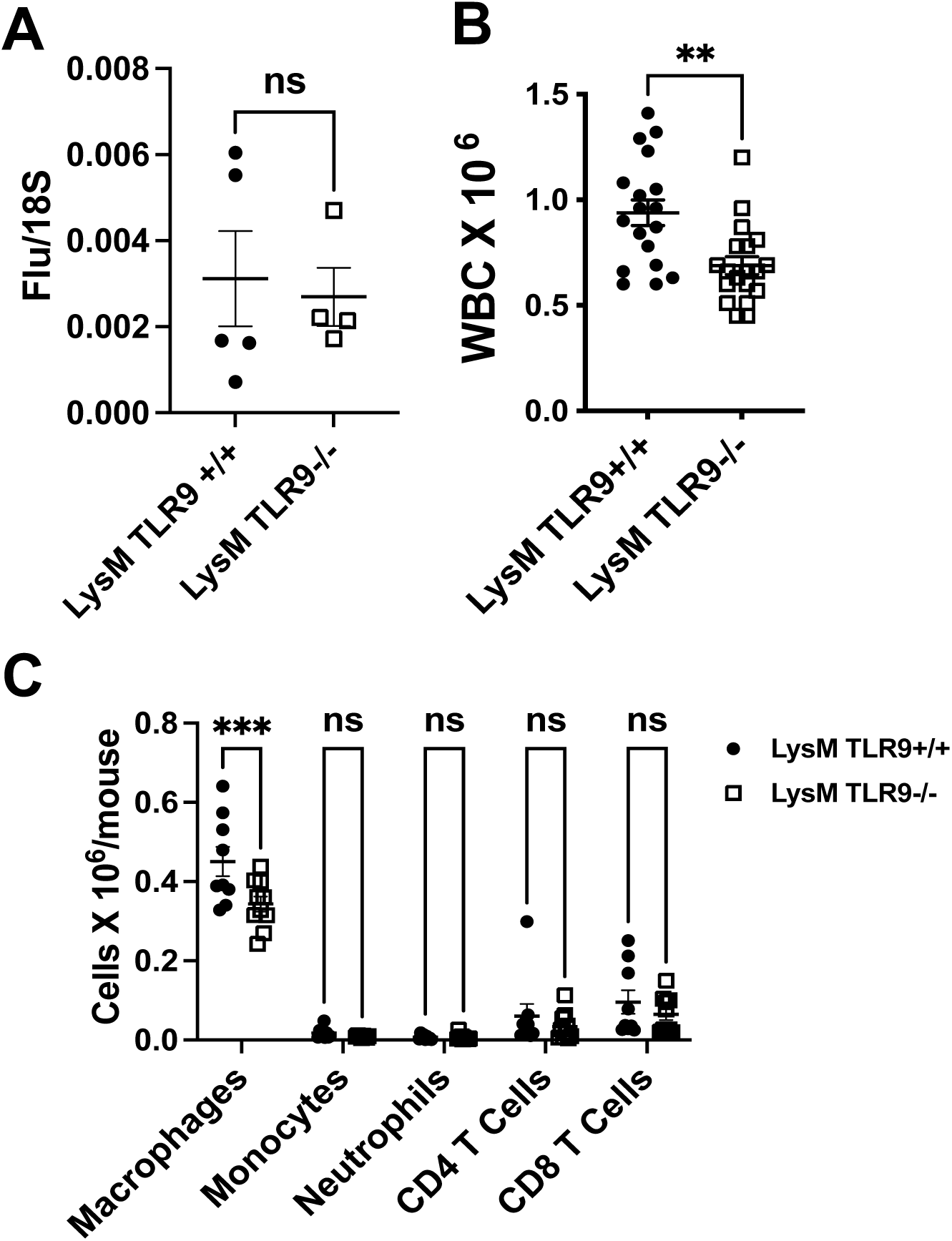
Myeloid specific TLR9 deficiency ameliorates lung inflammation without affecting viral clearance. LysM TLR9+/+ and LysM TLR9-/-were infected with influenza virus and euthanized on day 7 or day 14 to measure viral load and inflammation. Viral load was measured by qPCR on day 7 (A). Total WBCs were measured in the BAL samples harvested on day 14 (B) and specific cells were identified in a subset of experiments using flow cytometry (C). **, P < 0.01 using Mann-Whitney test and ***, P < 0.005 using two-way ANOVA.

**Supplemental Fig. 11.**
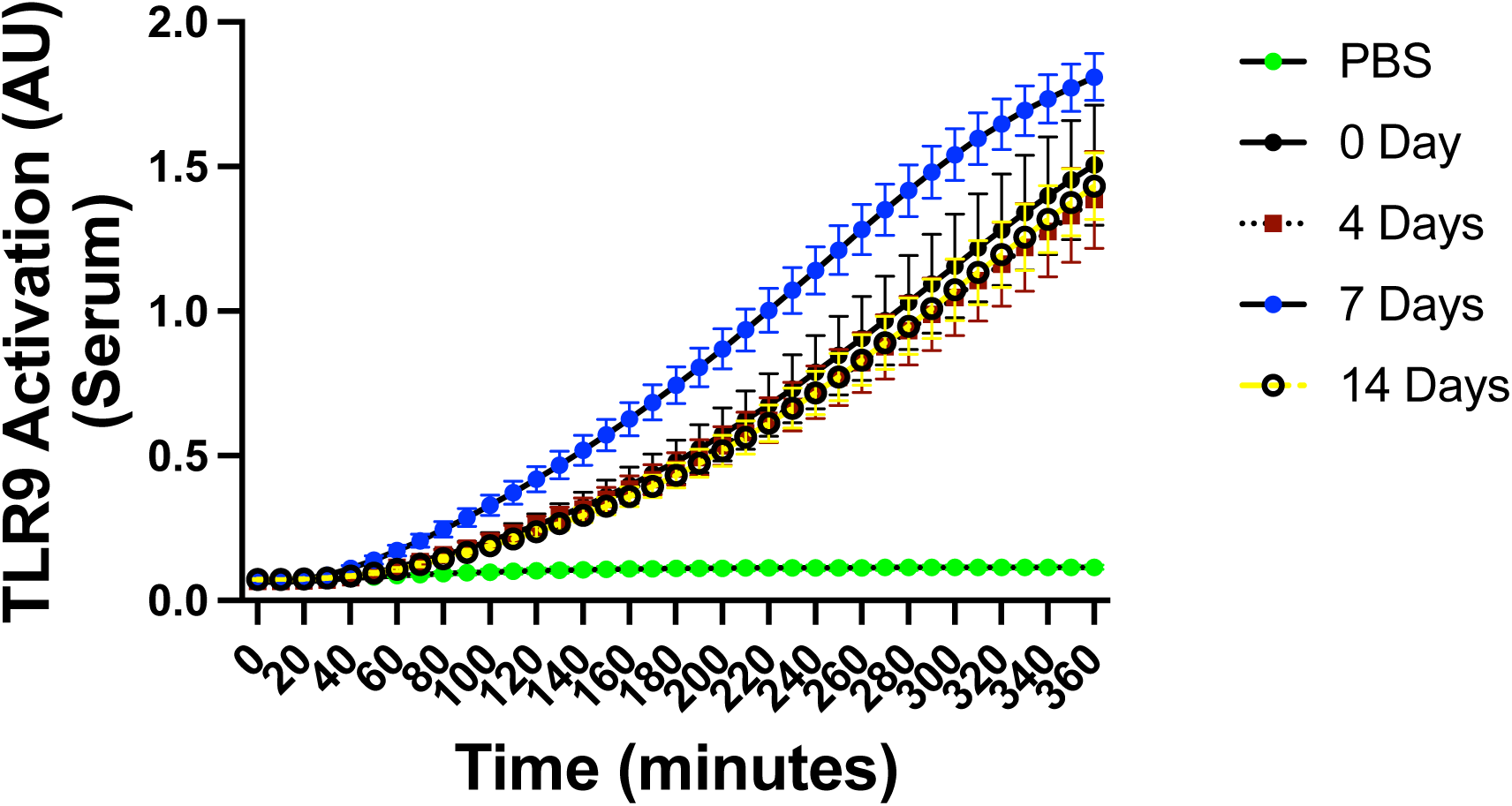
TLR9 activity was measured in mouse serum samples that were infected for 4, 7, or 14-days using mouse TLR9 reporter cell line. The data were measured using repeated measures for 6 hours. PBS alone served as a negative control. Day 7 values are statistically significant.

**Supplemental Table 1: Raw data that were used to generate all the figures used for this manuscript.**

